# Excitatory postsynaptic calcium transients at Aplysia sensory-motor neuron synapses allow for quantal examination of synaptic strength over multiple days in culture

**DOI:** 10.1101/2021.03.10.434766

**Authors:** Tyler W Dunn, Wayne S Sossin

**Author notes:** Corresponding Author. (WSS).

## Abstract

The ability to monitor changes in strength at individual synaptic contacts is required to test the hypothesis that specialized synapses maintain changes in synaptic strength that underlie memory. Measuring excitatory post-synaptic calcium transients through calcium permeable AMPA receptors is one way to monitor synaptic strength at individual synaptic contacts. Using a membrane targeted genetic calcium sensor, we demonstrate that one can measure synaptic events at individual synaptic contacts in Aplysia sensory-motor neuron synapses. These results show that synaptic strength is not evenly distributed between all contacts in these cultures, but dominated by multiquantal sites of synaptic contact. The probability, quantal size and quantal content can be measured over days at individual synaptic contacts using this technique. Surprisingly, most synaptic contacts were not found opposite presynaptic varicosities, but instead at areas of pre- and post-synaptic contact with no visible thickening of membranes. This technique shows promise in being able to address whether specialized synapses maintain synaptic strength underlying memory.

## Introduction

How long-term changes in neuronal systems underlie memory remains an important outstanding question in neuroscience. While much research has focused on changes that occur immediately after the stimulus, such as long-term potentiation, it is clear that what is often termed ‘consolidation’ or what perhaps is more correctly termed long-term memory formation, is not simply due to solidifying short-term changes (Sossin 2008). One attractive model for neuronal changes underlying long-term memory formation is the formation of new synapses (Bailey et al. 2015). While there are many correlative studies suggesting increased numbers of synapses after learning there are few systems where one can examine the number of synapses between identified neurons before or after a stimulus that generates long-term synaptic changes. This is important, because even with new technologies that have been able to document increased numbers of synapses between ensembles of neurons allocated to a memory compared to neurons not allocated to a memory (Choi et al. 2018), it is not clear whether the increased numbers of synapses was present before allocation and helped bias the allocation decision (Kim and Cho 2020), or whether synapse formation and stabilization occurred due to the stimulus. The ideal experiment would be to follow the connections before learning, document the increased number of synapses, and then determine whether the increased number of synapses was important for the increase in synaptic strength between these neurons thought to be important for memory maintenance.

The sensory-motor neuron synapse of Aplysia is an important model system that is well suited for this task. In this system one can monitor learning-related long-term changes in synaptic strength in a reduced preparation starting before the stimulation and lasting over one week after stimulation (Kandel 2001; Hu et al. 2017a). One of the striking observations in this preparation, is that similar to the animal, long-term changes in synaptic strength are accompanied by morphological changes, particularly an increase in presynaptic varicosities (Bailey and Chen 1988; Glanzman et al. 1990; Casadio et al. 1999; Bailey et al. 2015). These studies provide some of the strongest evidence that long-term increases in synaptic strength are supported by an increase in the number of synapses. In this system, blocking reconsolidation, or using pharmacological or dominant negative inhibitors of persistent protein kinases erase both sensitization and reduce long-term increases in synaptic strength back to baseline (Cai et al. 2011; Lee et al. 2012; Hu and Schacher 2015; Hu et al. 2017b). Surprisingly, while the number of varicosities decreased after erasure, the new varicosities formed after learning were not specifically removed (Chen et al. 2014). The lack of specificity for new varicosities has been interpreted as evidence that memories are not stored at synapses, but that instead, the nucleus retains a record of the baseline number of synapses and the modification of this number by nuclear changes would be the source of memory (Abraham et al. 2019). While the idea of non-synaptic memory has gained in popularity based on this finding (Tonegawa et al. 2015; Trettenbrein 2016), we have pointed out weaknesses in this argument (Sossin 2018).

One issue with the finding that erasure of synapses is not specific to new synapses, is that synapses in this study were measured by presynaptic varicosities, but these are only a surrogate for actual synapses. In the animal, only approximately 40% of sensory neuron presynaptic varicosities have active zones (Bailey and Chen 1983), and in cultures only approximately 30% of presynaptic varicosities actively release transmitter based on studies using synaptophluorin (Kim et al. 2003). Moreover, while increases in varicosities are usually highly correlated with increases in synaptic strength, there are exceptions. One day of training leads to sensitization without increases in varicosities (Wainwright et al. 2002); similarly if application of serotonin is restricted to the cell soma, 24 h increases in synaptic strength are observed in the absence of an increase in varicosities (Wainwright et al. 2002). Moreover, there are very few studies that systematically examine the percentage of synapses in this system that are not associated with varicosities.

To address this issue, better mechanisms for localizing synapses in sensory-motor neuron cultures are required. In vertebrates, many AMPA receptors are impermeable to calcium due to RNA editing of a critical glutamine in the AMPA receptor pore. However, in Aplysia there is no evidence for this editing in any AMPA receptor cDNA sequence (Greer et al. 2017) and invertebrate AMPA receptors without this modification are calcium permeable (Li et al. 2016). Thus, one can localize synapses in *Aplysia* using calcium-sensitive dyes that detect calcium transients in the post-synaptic cell that are locked to the action potential and these excitatory post-synaptic calcium transients (EPSCaTs) are blocked by AMPA receptor antagonists (Malkinson and Spira 2010). Using this technique approximately 56% of presynaptic varicosities adjoined post-synaptic calcium transients, but only 60% of synapses were at varicosities. However, these studies focused only on regions rich in varicosities and may have underestimated the number of synapses not at varicosities. Instead of calcium dyes used in this previous study, the use of sensitive genetically encoded calcium sensors that are membrane delimited can be used to detect synapse location in invertebrates (Akbergenova et al. 2018). Here, we present data using expression of a calcium sensor to localize synapses in unstimulated *Aplysia* sensory-motor neuron cultures. Using strict criteria to define EPSCaTs we find that most EPSCaTs are not found at varicosities. Even with the better localization of the calcium transient with a membrane-delimited calcium sensor, many EPSCaTs are made up of multiple quanta. EPScaTs were stable over days and this technique shows promise in being able to address the question of whether new synapses maintaining the increases in synaptic strength thought to be important for memory maintenance.

## Results

### Generation of a calcium sensor optimized for visualizing synaptic calcium transients

To address important issues of synapse specificity and whether new synapses formed by learning participate in the increases in synaptic strength (Sossin 2018) requires a technique that can both localize synapses over days in culture and measure the strength of individual synapses. Pioneering work using calcium indicator dyes showed that excitatory post-synaptic calcium transients (EPSCaTs), mediated mainly by calcium flowing through calcium-permeable AMPA receptors, can be used to localize and measure the strength at synapses in *Aplysia* sensory-motor neuron cultures (Malkinson and Spira 2010; Malkinson and Spira 2013). However, calcium dyes tend to compartmentalize over time (Paredes et al. 2008), making multiday use problematic. Moreover, their mobile cytoplasmic location results in substantial diffusion of the fluorescent calcium signal from its site of entry, limiting the fine resolution of synapse location. Genetically encoded calcium sensors, such as GCaMP (Nakai et al. 2001) can solve these issues. Adding protein determinants that limit the GCaMP to membranes can decrease the spread of the synaptic calcium fluorescence signal, as calcium entering across the plasma membrane would most likely interact with the membrane delimited GCaMP subpopulation at the site of entry, the synapse, have minimal diffusion while fluorescent as the calcium bound GCaMP is membrane bound, and then release calcium into the cytosol with a much greater chance of subsequent buffering through other, non-fluorescent mobile and non-mobile calcium buffers (Shigetomi et al. 2010). We limited GCaMP6s to membranes by fusing it to the Aplysia Src N-terminal that contains myristylation and palmitylation signals (src-GCamP6s), similar to previous membrane delimited versions of GCaMP (Shigetomi et al. 2010; Akbergenova et al. 2018). We believe this approach is better than using a transmembrane domain due to the lack of a requirement for passage through the secretory system, often leading to accumulation in the ER, as we have observed for other tagged transmembrane proteins expressed in Aplysia sensory neurons (Nagakura et al. 2008; Farah et al. 2019).

### Src-GCAMP6s is effective at measuring action potential evoked calcium transients

Single *Aplysia* sensory neurons were cultured with single LFS motor neurons for three days. On the third day, nuclear injection of plasmids containing RFP into the presynaptic sensory neuron and src-GCaMP6s into the postsynaptic motor neuron was carried out and the cultures left a further day to allow for plasmid expression (Fig. 1A). Intracellular sharp electrodes were used to control and measure membrane potential of both the presynaptic and postsynaptic neurons. Image acquisition (25ms exposure, ~60ms frame rate) was synchronized to the stimulus pulse to standardize data acquisition of evoked synaptic transmission and allow for automatic processing of the large data set associated with each experiment. We acquired 9 images to serve as a baseline fluorescence signal, evoked a presynaptic action potential during frame 10, and continued imaging for 35 more frames. Subtraction of the frame preceding the action potential (frame 9) from the frame immediately following the action potential (frame 11) produces an image of the calcium transient occurring during the given stimulus (Fig. 1B).

**Figure 1.**
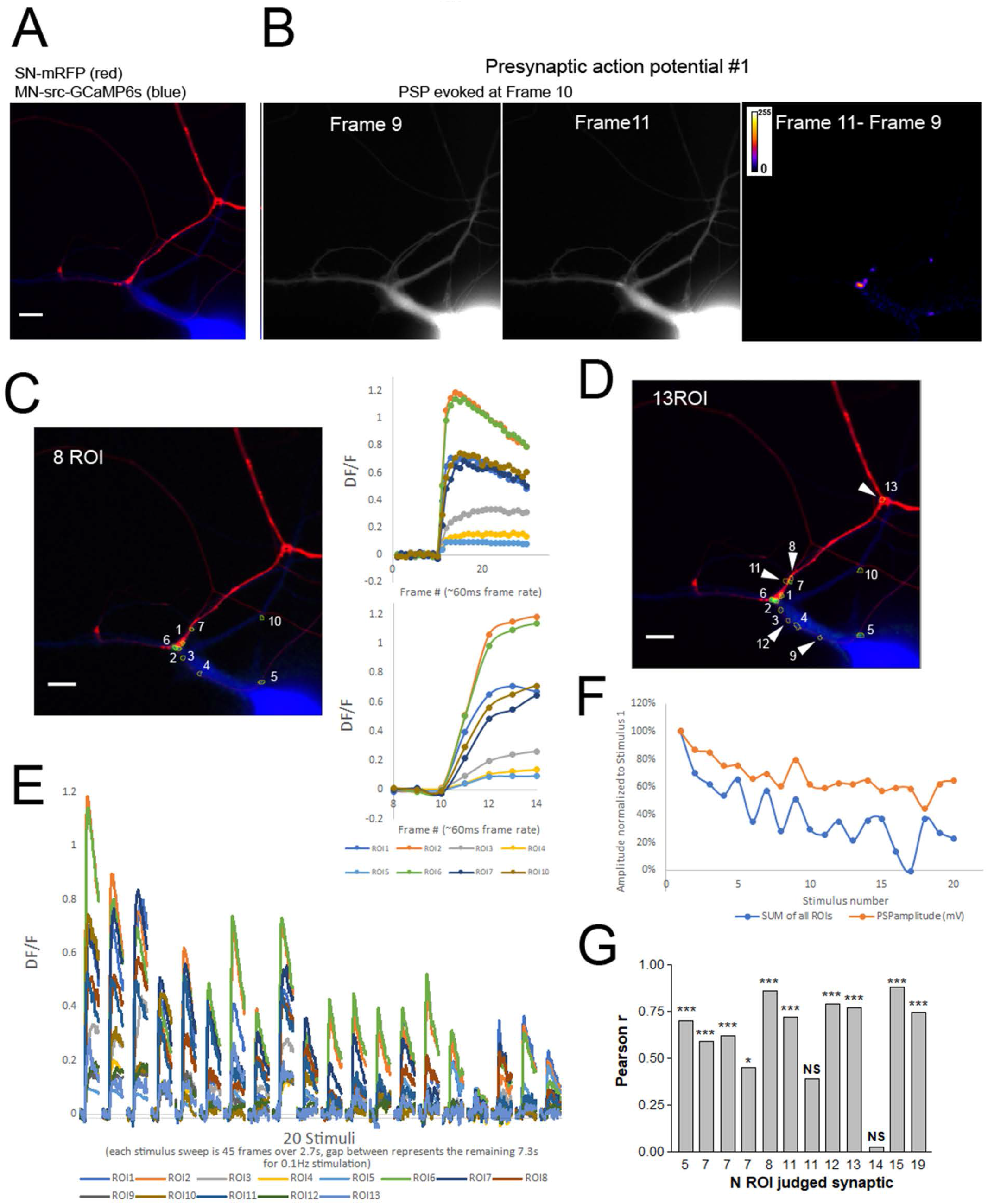
Evoked synaptic transmission at the *Aplysia* sensory to motor neuron synapses measured with postsynaptic src-GCaMP6s calcium transients. A) Image of synaptic pair, with the presynpatic neuron expressing RFP (red) and the postsynaptic neuron expressing src-GCaMP6s (blue). B) With 45 frames taken, an action potential is evoked during the tenth frame. Subtracting the src-GCaMP6s signal from frame 9 (taken immediately preceding the action potential), from the src-GCaMP6s signal from frame 11 (taken immediately following the action potential) highlights the src-GCaMP6s transient in the resulting produced image (right panel, intensity presented in a ‘fire’ lookup table with the calibration bar for the calculation results of the image subtraction in arbitrary units). Scale bar is 20μm. C) Using the same image in A for the presynaptic RFP and postsynaptic src-GCaMP6s (blue), the frame 11-9 image subtraction image for a single stimulus (Stimulus #1) is in the green channel and is used to find ROIs that are labeled. Intensity at each ROI is normalized into a DF/Fo using the average of frame 1-9 as Fo. The transients can be seen to have variable amplitude and various time courses (upper right). The rise rate of the transients can be seem more clearly just focusing on frames 9-14 (lower right). D) To gather additional data about the synaptic connections examined, 20 stimuli are delivered at 0.1Hz. Red and blue channels are as in A, however the green channel is now the summation of the frame11-frame9 images for all twenty stimuli. Since not all synapses participate in the first stimulus, additional ROI are identified (5 new ROIs shown by arrows in the example shown). E) The normalized transients at each of the ROIs shown in C are overlayed to show the variation in amplitude between ROIs and from stimulus to stimulus. F) PSP amplitude undergoes characteristic homosynaptic depression with these twenty successive presynaptic action potentials, as does the intensity of the summed DF/F values measured at all the ROIs with strong correlation between the two measures (Pearson r 0.8265, P<0.000001). G) Pearson r values from synaptic connections examined with 5 or more ROIs within the field of view (number of ROI in the trial as x-axis label). Only synapses with a Pearson r value that produces a P value P<0.05 will be further examined, therefore trials with less than 5 ROI or a P>0.05 would not be used in analysis (** P<0.01, *** P<0.001). The two trials removed for having a non-significant correlation between PSP amplitude and the evoked src-GCaMP6s transient amplitude are shown.

To identify and analyze synaptic transmission using the postsynaptic calcium transient, regions of interest (ROIs) were selected as described in the methods. Multiple ROIs with varying magnitudes of fluorescent changes are observed dispersed throughout the field of view after a single stimulus (Fig. 1C). The peak of the transients tends to occur much later than frame 11, however, as frame 12 is acquired 80-110ms following the peak of the PSP, and therefore the end of the synaptic current, the source of the calcium leading to the further increases in DF/F beyond frame 11 are likely from sources other than entry through synaptic glutamate receptors, such as L-type calcium channels (Malkinson and Spira 2010) and thus, only data from frame 11 is used for measurements of synaptic strength. In order to maximize the descriptive data from every sensory to motor neuron pair examined, twenty stimuli were evoked per trial at 0.1Hz. Summation of all twenty calcium transient images (Frame 11-Frame 9 for each stimulus) better describes the entire synapse within a field of view, often revealing sites that did not participate in the first stimulus (Fig.1D). The number of new ROIs identified with subsequent stimuli reduce so that only 2.0±1.0% (n=10 synaptic connections) of the ROIs first participate in one of the last five stimuli, suggesting that few additional sites would be found with more stimuli. However, if there are very low probability synapses (probability of release near or less than 0.05), these synapses will be undercounted. Moreover, as described below, using twenty stimuli allows for better resolution of the individual ROIs and a far greater data set for more accurate analysis of the quantal nature of the EPSCaTs. Following generation of the full ROI set for a particular synaptic pair, fluorescence intensity is measured for all ROIs over the twenty evoked synaptic stimuli, converting fluorescence intensity into DF/F values (Fig.1E). The transients at each ROI over the twenty-stimulus trial are independent and variable in amplitude, consistent with transmitter release from independent synapses synchronized by the presynaptic action potential (Fig. 1E). Comparison of the normalized PSP amplitude to the normalized EPSCaT amplitude for each of the twenty stimuli shows a strong correlation between these two measurements (Fig. 1F). Most trials with five or more ROIs in the field of view show a strong correlation between these two independent measures of synaptic transmission (Fig. 1G). However, in some circumstances, either due to strong and widespread burst of non-synaptic calcium transients or to a high proportion of the synapses being outside the field of view, this correlation was not significant and these trials were not used in the analysis (Fig. 1G)

In order to maximize the chance of seeing all synapses within the field of view, we increased the extracellular calcium to magnesium ratio as previously done for measuring EPSCaTs with calcium indicator dyes (Malkinson and Spira 2010; Malkinson and Spira 2013). To determine the extent to which this affected synaptic transmission, we measured the effect of this high calcium recording saline on the amplitude of the PSP. Since the *Aplysia* sensory to motor neuron synapses undergo homosynaptic depression, we measured the change in PSP amplitude going from both normal to high calcium saline and from high to normal calcium saline. The difference in the change in PSP amplitude between these two measured equal to the change in PSP amplitude with high calcium saline and was found to be about ~15% (Fig. 2).

**Figure 2.**
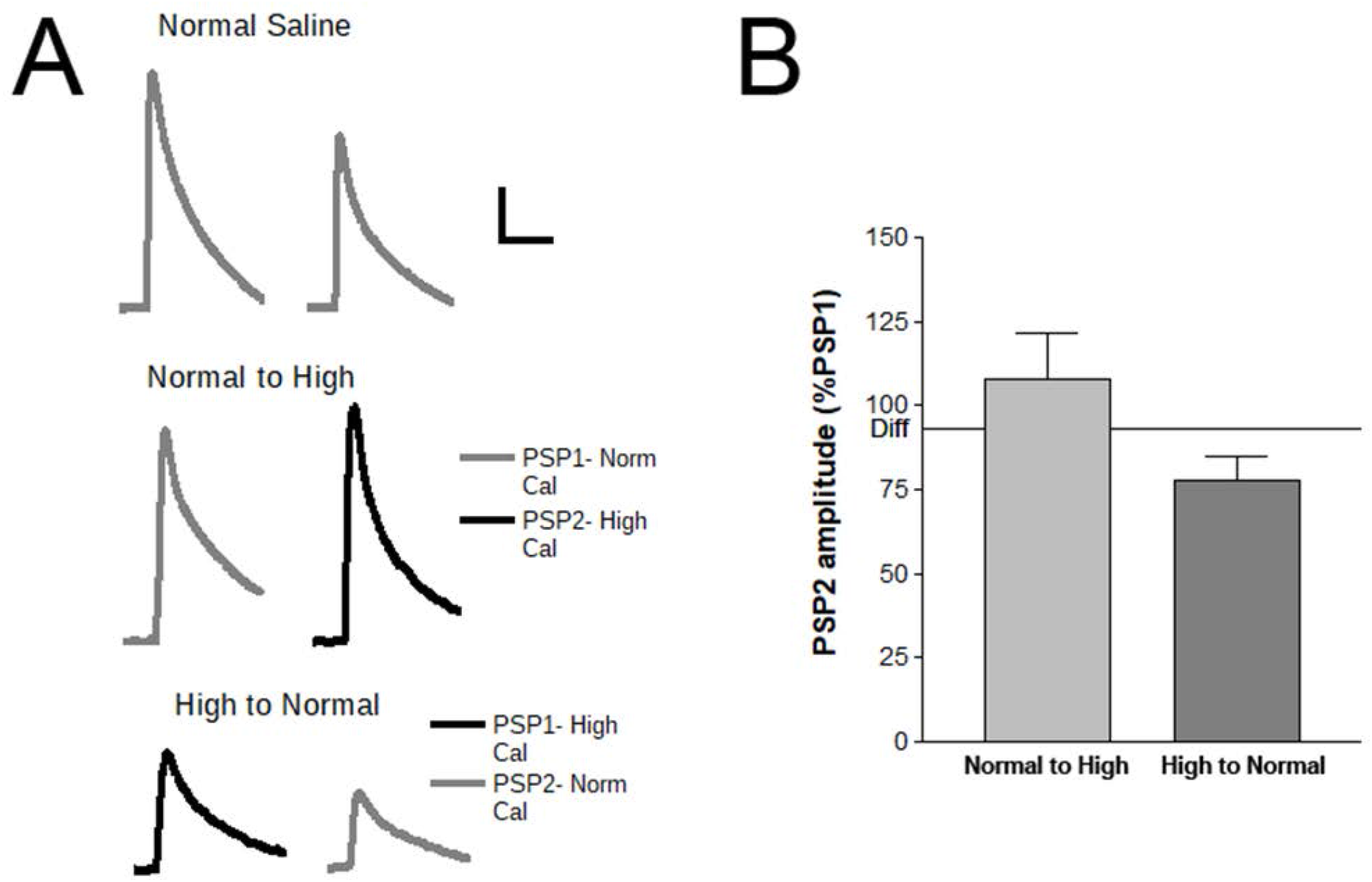
High calcium saline on PSP amplitude. A) Representative PSP traces. Top traces, PSP1 and PSP2 in normal saline (highlighting depression by HSD). Middle traces, with PSP1 in normal saline, PSP2 evoked following 30mL perfusion with high calcium saline. Lower traces, with PSP1 in high calcium saline and PSP2 evoked after perfusion of 30mL of normal saline. Scale bars are 10mV/50ms. B) Summary of the average PSP amplitude change going from normal to high calcium saline (n=6 synaptic connections) and from high calcium to normal calcium saline (n=7 synaptic connections). SEM. Since all experiments will undergo synaptic depression and this high calcium buffer does not affect the amount of depression (Malkinson and Spira 2013), the effect of high calcium can be simply calculated by the average of both amplitude changes. This is marked with a bar and is equal to a change in PSP amplitude with high calcium saline at ~15%.

### Developing criteria for separating EPSCaTs from other calcium transients

Src-GCaMP6s transients can occur frequently in the postsynaptic neuron without any correlation to synaptic membrane potential changes either evoked or spontaneous. These transients have variable time courses and can appear similar to the synaptic transients (Fig. 3A). While the size of the ROI tends to be much larger with the non-synaptic transients than those of the synaptic ROIs, the time course of some of these events can appear similar to the synaptic events. In some instances, these transients occur at or near enough to selected ROIs to create coincidental false positive transients during the action potential stimulus. To reduce the impact of these transients, that are not directly involved in synaptic transmission, we added an additional criteria at each ROI for inclusion of each transient in subsequent analysis. If a transient initiates prior to frame 10, when the presynaptic action potential occurs, the transient is considered a calcium transient unrelated to direct synaptic transmission. As the non-synaptic calcium transients have variable rates of occurrence within and from trial to trial, the level of interference in measuring synaptic transmission also varies. The number of events that initiate prior to the stimulus at a chosen ROI range from 0 up to a rate of 4.3%, averaging 0.9±0.2% (data from 18 synaptic connections). From this we can estimate that a similar percentage of non-synaptic calcium transients are also occurring at synaptic ROI during the period when synaptic transients are measured (frame 10 and 11 initiation) producing false positives that slip through this filter. Examples of non-synaptic transients that interfere with EPSCaT measurement are shown in Fig. 3B and removed from subsequent analysis. Both trials removed for having a non-significant correlation between PSP and EPSCaT amplitude (Fig. 1G) had a high number of these transients (3.6 and 4.3% respectively).

**Figure 3.**
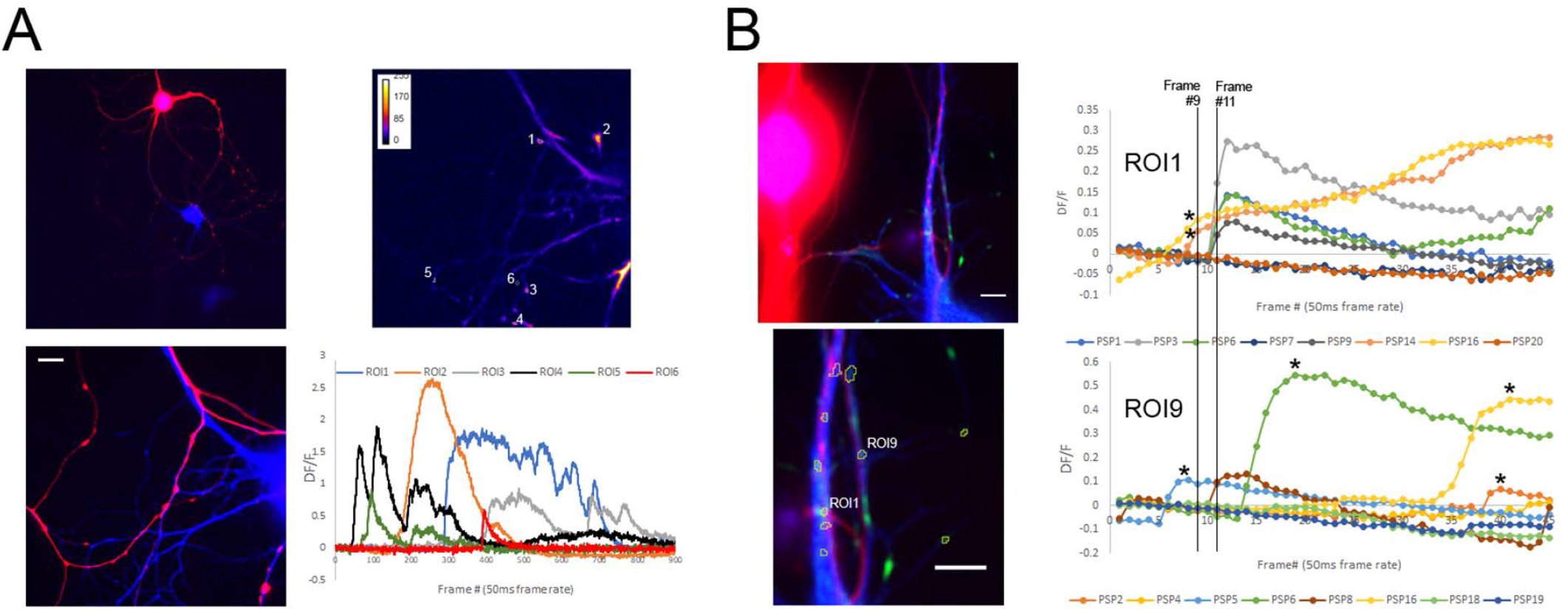
Non-synaptic calcium transients. A) Representative image of sensory neuron expressing RFP and the postsynaptic neuron expressing src-GCaMP6s, 10x image above 40x image (scale bar is 20μm). Upper right image is a compressed image of the maximum pixel value over a 900 frame acquisition period subtracted from the first, baseline frame with intensity presented in a fire look-up table, highlighting the location and variable size of the non-synaptic calcium transients (presynaptic neuron was kept quiescent with hyperpolarization and the postsynaptic neuron monitored for any spontaneous mini synaptic transmission during the imaging period). The fluorescence intensity of the ROIs selected in this image are graphed below showing the variable time-courses of these non-synaptic calcium transients, using the first 10 frames to measure Fo for each ROI. B) A single trial demonstrating the potential for non-synaptic events to appear synaptic. Two synaptic ROIs from the synaptic pair shown on the left (magnified in on lower image) chosen for the high number of non-synaptic events occurring during the action potential evoked synaptic transmission measurements marked with *. Only some of the twenty PSP sweeps are shown to highlight the synaptic transients (that initiate at frame 11) and the events not directly measuring the synaptic current either through early or late initiation (*). Scale bars are 20μm.

### Dissociation of independent ROIs within a cluster with repeated stimuli

Examination of the individual synaptic ROIs reveals a homogeneous area distribution and apparent quantal fluctuations in the action potential evoked calcium transients measured with src-GCaMP6s. The representative trial shown in Fig. 4A-C displays how the synaptic calcium transients of three closely spaced ROIs are independent in participation in the series of stimuli and can appear to fluctuate from failures to transients of consistent amplitude (marked with dotted line). However, transients often appear to be many times larger than the minimum events, as with PSP6 at ROI3 in the example (Fig. 4A-C). Whether this apparent multi-quantal synaptic transmission is the result of intra-site multivesicular release, or closely spaced synapses is unclear and cannot be resolved solely by EPSCaT measurements. To resolve the independent ROIs within the large clusters of synapses sometimes observed and highlighted at the synaptic connection shown in Fig. 4D, ROIs are chosen from PSP20 to PSP1. As displayed in the montage of twelve PSPs in Fig. 4E, the independent participation of the synapses within a cluster allows for refining of the ROI set, such independence of sites is visually apparent if each PSP is given a different color LUT and with all colors combined in an overlay, the color combinations can highlight the independent ROIs participating in the cluster (Fig.4F). By selecting and refining all the ROIs using the data from all the stimuli allows separation of many of the independent ROIs within a cluster, but we may not have sufficient resolution to fully achieve this. Examination of the mean ROI area (9.0μm±3.5SD, n=110) over many synaptic connections finds a homogeneous distribution of ROI areas. The distribution was compared to a Gaussian distribution with a Runs test with no significant deviation from the model (P=0.7731), however, some events do appear about twice the size of the mean suggesting some ROIs may still be tight clusters of multiple independent synapses (data from 10 synaptic connections).

**Figure 4.**
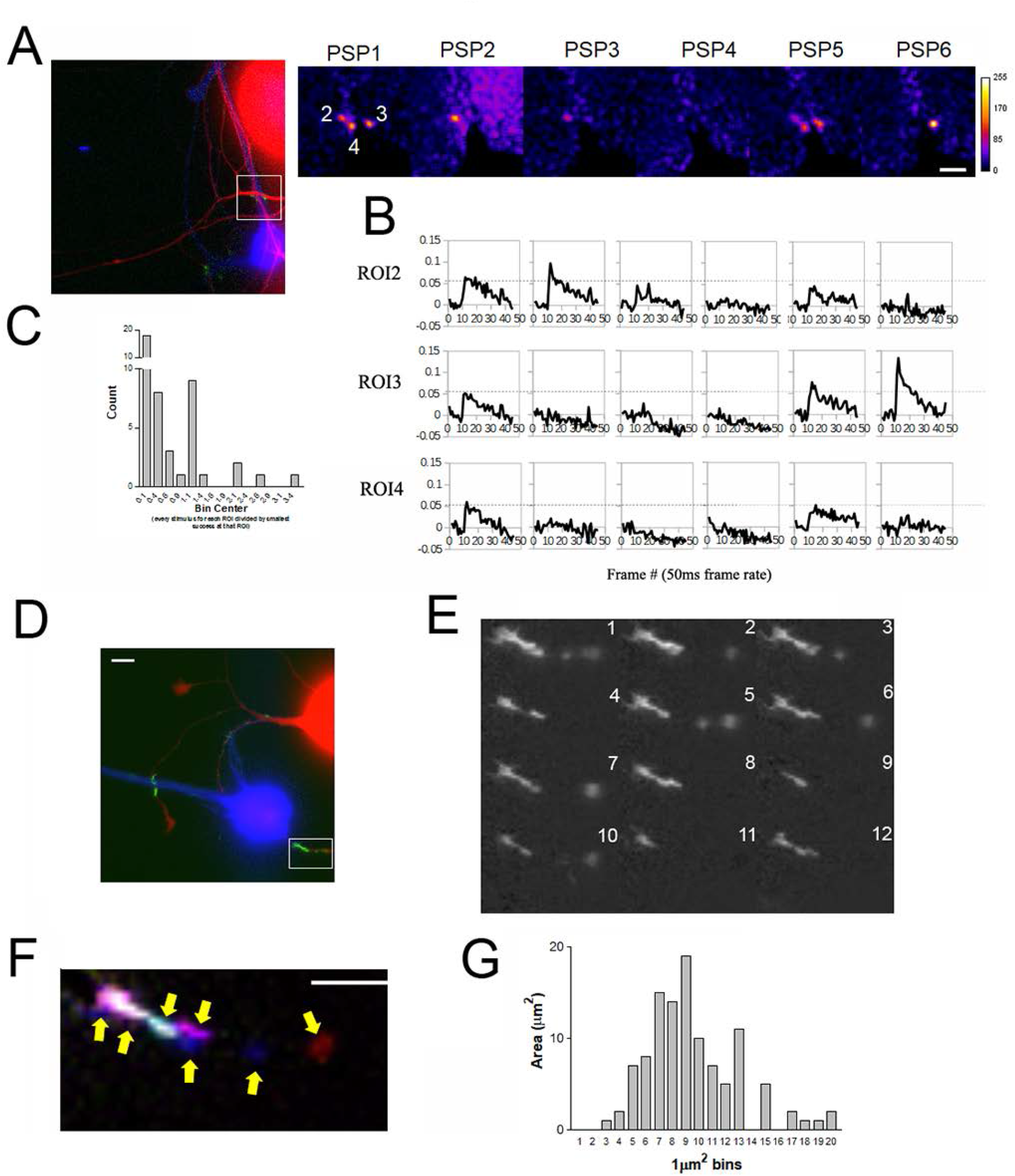
Quantal fluctuations and clusters of independent synapses. A) Left image is presynaptic RFP, postsynaptic src-GCaMP6s (blue), sum of the EPSCaTs (frame 11-frame 9) for the 20 stimuli (green), with the location of the magnified right images PSP1 to PSP6 located with the white box. The three ROIs located in this sub-region show apparent quantal fluctuations with successive stimuli, visually apparent with the single stimulus EPSCaT images (frame 11 – frame 9 image subtraction generated) colored with a fire look-up table to enhance the contrast (units of subtraction are arbitrary with the color palette calibration bar on the far right, scale bar is 10μm). B) The fluorescence intensity data from the same ROIs and stimuli shown in A, collected over the 45 frames converted to DF/F values. The dotted lines are for reference comparison of amplitudes within a trial. C) Frequency histogram of frame 11 DF/F values for all twenty stimuli at the seven ROI found at the synaptic connection shown in A (including failures of any evoked transients) normalized to the minimum observed synaptic transient at each ROI. Most events bin into the noise band and represent synaptic failures. The second peak occurs around 1 as many synaptic transients are similar in amplitude, however, events are also seen at multiples of this value. D) Presynaptic RFP (red), postsynaptic src-GCaMP6s (blue), and the DF/Fo (green) at a representative synaptic connection with a cluster of independent synapses. E) Montage of 12 successive stimuli evoked EPSCaTs at the cluster of synapses magnified from the box in D with the number of the stimulus in the right top of each image. Fluctuations in transmission at the independent sites in the cluster are apparent as sites variably participate in each stimulus. F) Pseudo-coloring each frame from D with a different color and merging them in an overlay reveals a tight cluster of independent sites marked with arrows (scale bar is 10μm). G) Frequency distribution of ROI areas measured from 10 synaptic connections with at least 5 ROIs binned in 1μm2 bins.

### Large EPSCaTs represent multiquantal transmission

Even with this separation of the clustered ROIs, many ROIs still show transients that are many multiples of the smallest transients observed at the ROI (Fig. 5A). To further examine the amplitude variance of the postsynaptic src-GCaMP6s transients at the individual ROIs (Fig.5A) we designed a set of logic steps to automatically sort events and estimate quantal size at individual ROIs, assuming the variation is largely the result of quantal fluctuations. At each ROI, quantal size was determined by the minimum synaptic transient amplitudes and the variance of all synaptic transients at the ROI (Fig. 5B). The ROI minimum, median, and maximum event amplitudes are found, and cluster analysis used to group all events at the ROI into one of these three groups by proximity. If the maximum and minimum clusters are within 50%, all events would be averaged to find quantal size at the ROI, and such an ROI would be judged uniquantal as all events are of a consistent amplitude. At ROIs with more variation, quantal size is found from the average of the median and minimum clusters if these clusters are within 50%, if not quantal size is found from the average of the minimum cluster (Fig. 5CD). Once quantal size is estimated, a quantal content estimate can be made at each ROI for every stimulus, generating a quantal content heat map describing the synaptic connection within the field of view (Fig. 5E). From this representation of the data the success/failure probability of participation, the relative contribution of each ROI to the total quantal content, and number of ROIs participating in each stimulus can be readily derived (Fig. 5E). The sum of the quantal content estimates at all ROIs and the number of ROIs participating in each stimulus reduced at similar rates over the twenty stimulus trials (normalized to the first stimulus) similar to the rate of PSP amplitude during depression (Fig. 5F). As an independent measure of quantal size, we electrophysiologically measured miniature PSPs (mPSP) prior to imaging action potential evoked transmission. If our assumption that the high variance of synaptic transient amplitude seen at some ROIs is largely the result of quantal fluctuations and if our above quantal size estimates are accurate, the reduction in amplitude variance of the src-GCaMP6s transients judged uniquantal (ex. all the ones in Fig.5E) should better match the variance of mPSP amplitudes. When all measurements are normalized to the minimum at each synaptic connection we find that the uniquantal estimates have a similar distribution and variance to the independently measured mPSPs (Fig. 5G). The high variance observed when examining all transients is much greater than both the events judged uniquantal and mPSP amplitudes, indicating that the source of most the variance at the ROIs with high transient variance is multiquantal synaptic transmission at the ROI and that our method of estimating quantal size appears accurate. Thus, at each ROI, quantal size, quantal content, and the probability of participation in action potential evoked release can be estimated for each ROI at a synaptic connection.

**Figure 5.**
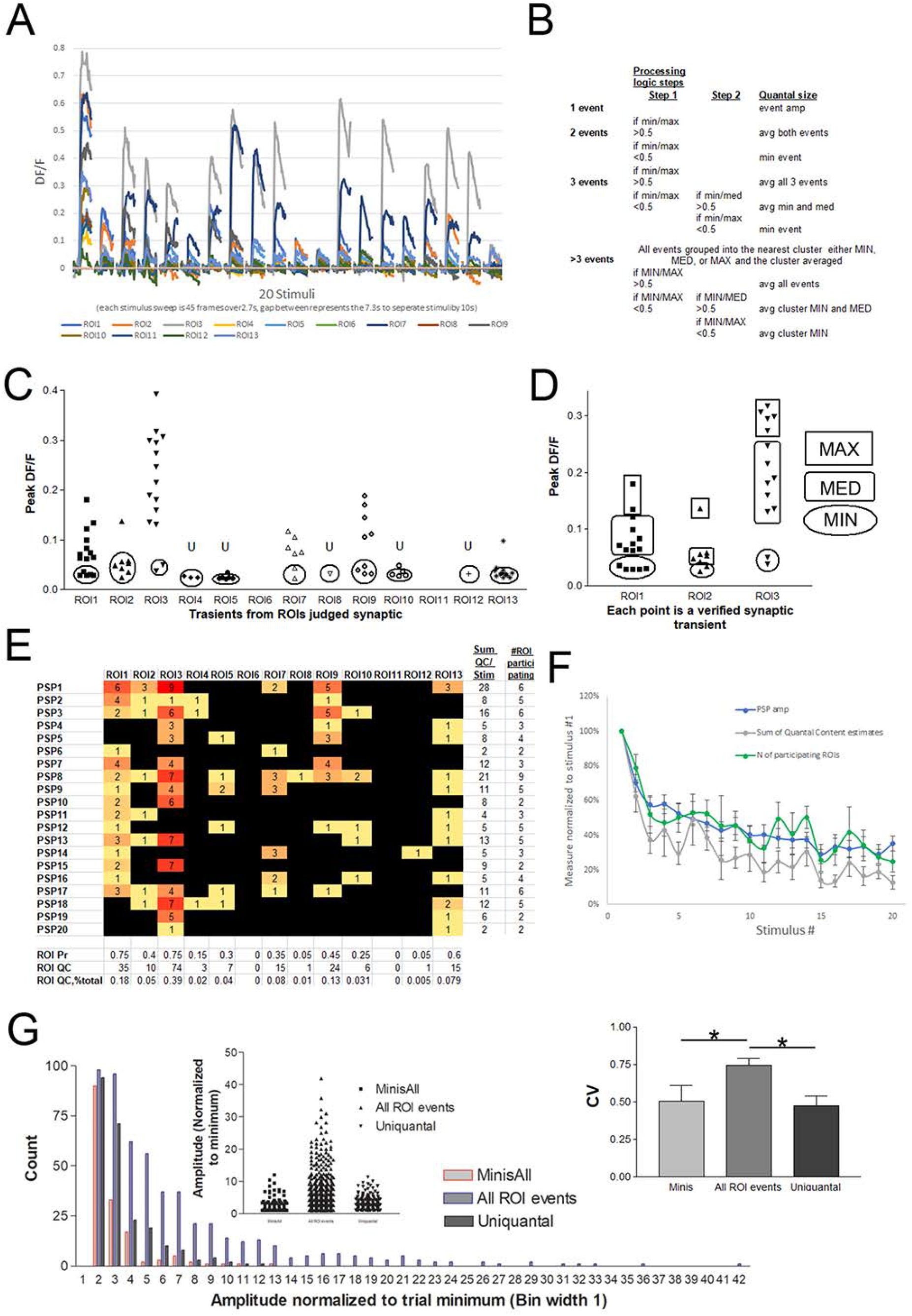
Deriving the parameters of synaptic transmission from the src-GCaMP6s signal at the individual ROIs. A) Twenty successive stimuli generate src-GCaMP6s transients of variable amplitude, shown are the transients for a representative synaptic connection with 13 ROIs selected as in figure 1. B) Written description of the processing steps taken to determine quantal size at each ROI from the amplitude of the src-GCaMP6s transients to the twenty stimuli. Sorting and quantal size calculation was performed with excel spreadsheets. C) Peak amplitudes of transients judged synaptic at each ROI, with the group of transients averaged to determine quantal size as in B. Note that two ROIs fail to produce a single successful transient, either because no transient crossed the signal to noise ratio and/or were judged non-synaptic as in figure 3. The group of transients judged uniquantal at each ROI are circled and the ROIs judged uniquantal marked with a U. D) Highlighting the cluster analysis grouping outcomes for the first three ROIs from C, shows how data with differing variances are grouped. In the example, the minimum cluster is averaged for quantal size for ROI 1 and ROI3, whereas the minimum and median clusters are averaged in ROI2 based on the rules of B. E) Following automatic determination of quantal size, ROI data can be converted to a quantal content (QC) value for each stimulus, and a heat map generated to visually describe the synaptic connection as a whole, over all ROIs for every stimulus. From this, the total quantal content over the twenty PSPs (ROI QC), the probability of observing a synaptic transient at the ROI (ROI Pr, as proportion of successful synaptic transients observed), and the % the ROIs QC contributes to total QC at the synaptic connection (ROI QC % total) can be found for each ROI. Furthermore, the summed QC and number of participating ROIs can be found for each stimulus. F) Summary and comparison of the change in PSP amplitude to the sum of the quantal content estimates at all ROIs and the number of participating ROIs over the twenty stimuli recording period derived as in E. (Data from 10 synaptic connections). G) Comparison of the amplitude distribution of spontaneous mPSPs (minis) to the amplitude distribution of all synaptic ROI transients and to the subset of synaptic transients judged uniquantal as in B-E. mPSPs were recorded prior to stimulation and only synaptic connections with more than ten mPSPs recorded were used for analysis here. Normalization of all data within a trial to the minimum value for the measure at each ROI, allows comparing data between measures and from the six synaptic connections examined. The skew of the amplitude distribution of the synaptic ROI transients judged to be uniquantal was similar to the skew of the mini PSP amplitudes, as summarized with the coefficient of variance (CV) in the normalized amplitudes. The normalized amplitude distribution of all synaptic transients measured at the ROIs was much more skewed than both the electrophysiologically measured mPSPs and the synaptic transients judged to be uniquantal with a significant increase in the CV (comparing the CV of the normalized mPSP amplitude, all synaptic ROI transients, and only the uniquantal synaptic transients with a repeated measures AVOVA, * P<0.05, n= 6 synaptic connections).

### Most EPSCaTs are not associated with presynaptic varicosities in contact with the postsynaptic neuron

To examine whether most EPSCaTs occur apposed to presynaptic varicosities, mRFP expressed in the sensory neuron was used to blindly identify varicosities according to criteria outlined previously for sensory neuron-LFS synapses (Chen et al. 2014) (Fig. 6A). Using the methods described above to select ROIs that identify the location of the synaptic elements contributing within the field of view, we find that while many varicosities contain synapses, the majority do not (Fig. 6B). Probably more importantly, most EPSCaTs are not apposed to presynaptic varicosities, but instead are localized to regions where the presynaptic neuron grows on the postsynaptic neuron without obvious thickening on either side (Fig. 6B). Thus, in the sensory-motor neuron cultures used in this study, the presynaptic varicosities are not good indicators of the location of synapses defined by the EPSCaTs. As some ROIs appear to be multiquantal, we further sorted the data to examine what type of ROI tends to occur at varicosities. On average ~47% of the ROI in this data set were judged multiquantal (3 or more quantal content) while a slightly lower percentage of ROI at varicosities were judged to be multiquantal at ~31%, indicating no trend for the occurrence of multiquantal ROI specifically at varicosities.

**Figure 6.**
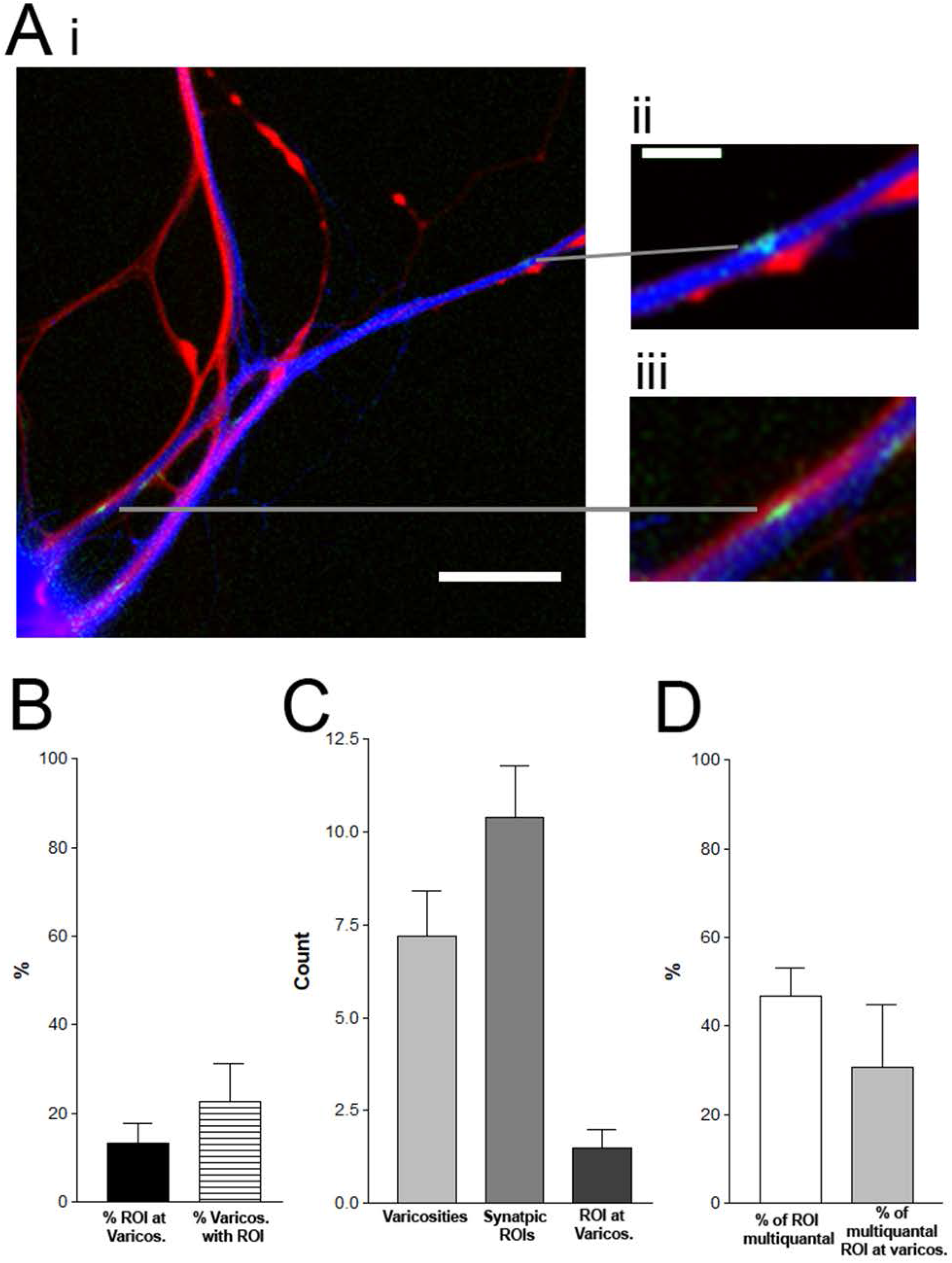
Synapse localization relative to presynaptic varicosities. A) Representative synaptic pair with the sensory neuron expressing RFP (red), the postsynaptic neuron expressing src-GCaMP6s (blue), and the location of the synapses visualized as in figure 1 (green). Presynaptic varicosities in contact with the postsynaptic neuron are identified by a blind judge and then compared to the location of the synaptic ROIs. An example of a synapse at a presynaptic varicosity (ii) and a synapse not at a presynaptic varicosity (iii) are shown in the insets. Scale bars 40μm in i, 10μm in ii, same scale in iii. B) Summary data from 10 synaptic connections, displaying the percentage of synapses (ROIs with synaptic transients) at presynaptic varicosities and the percentage of presynaptic varicosities with identified synaptic ROIs. C) The average number of varicosities identified by the judges and the average number of synaptic ROIs from the data set summarized in B. D) The percentage of all ROIs that appear multiquantal (3 or more multiples of the quantal size estimate at the ROI) and the percentage of ROIs at varicosities that appear multiquantal from the data set.

### Postsynaptic src-GCaMP6s can be used to track synapses over multiple days

To determine whether the described methods of measuring both the strength and spatial distribution of the synapse within a particular field of view can be used to track a synapse over multiple days we tracked synaptic pairs over two days (Fig.7). Some ROIs are clearly tracked over both days, having specific spatial localization in isolation and proximity to pre and postsynaptic image landmarks (Fig. 7A). The quantal content heat maps generated for each day as in figure 5 can be reorganized, so that the ROIs blindly judged to be the same between days, are reorganized into a single column. Some ROIs appear to move from day to day, or alternatively, sites are lost and new sites arise (new columns) (Fig. 7B). Some sites are close to the image boundaries so that their appearance or disappearance may reflect entry or exit of the filed of view, such as ROIs 17 and 20 in the example. Other ROIs are not faithfully tracked over both days, suggesting there are new synapses created and old synapses removed. Alternatively, the ROIs that appear may simple be dispersed from large clusters unresolvable on the previous day and conversely the loss of ROIs the clustering of multiple independent synapses, as suggested by the multiquantal nature of the ROIs. Anecdotally to this, ROI19 and ROI16 that disappear on day 2 are neighboring ROI9 and ROI13 respectively, two ROIs that appear highly multiquantal and have increased quantal content on the second day of imaging suggesting the possibility of sites merging over time. Importantly, neither the presence of src-GCamP6s over multiple days nor the use of the high calcium saline for the measurement of the EPSCaTs has any toxic effect on the synapse, since PSP amplitude is constant over both days (Fig. 7C). We found no significant change in the measure of quantal size and the sum of quantal content (both measured as described in figure 4), or the number of ROIs with synaptic transients (Fig. 7C), suggesting that this technique can be used to measure changes in synaptic strength over multiple days in culture.

**Figure 7.**
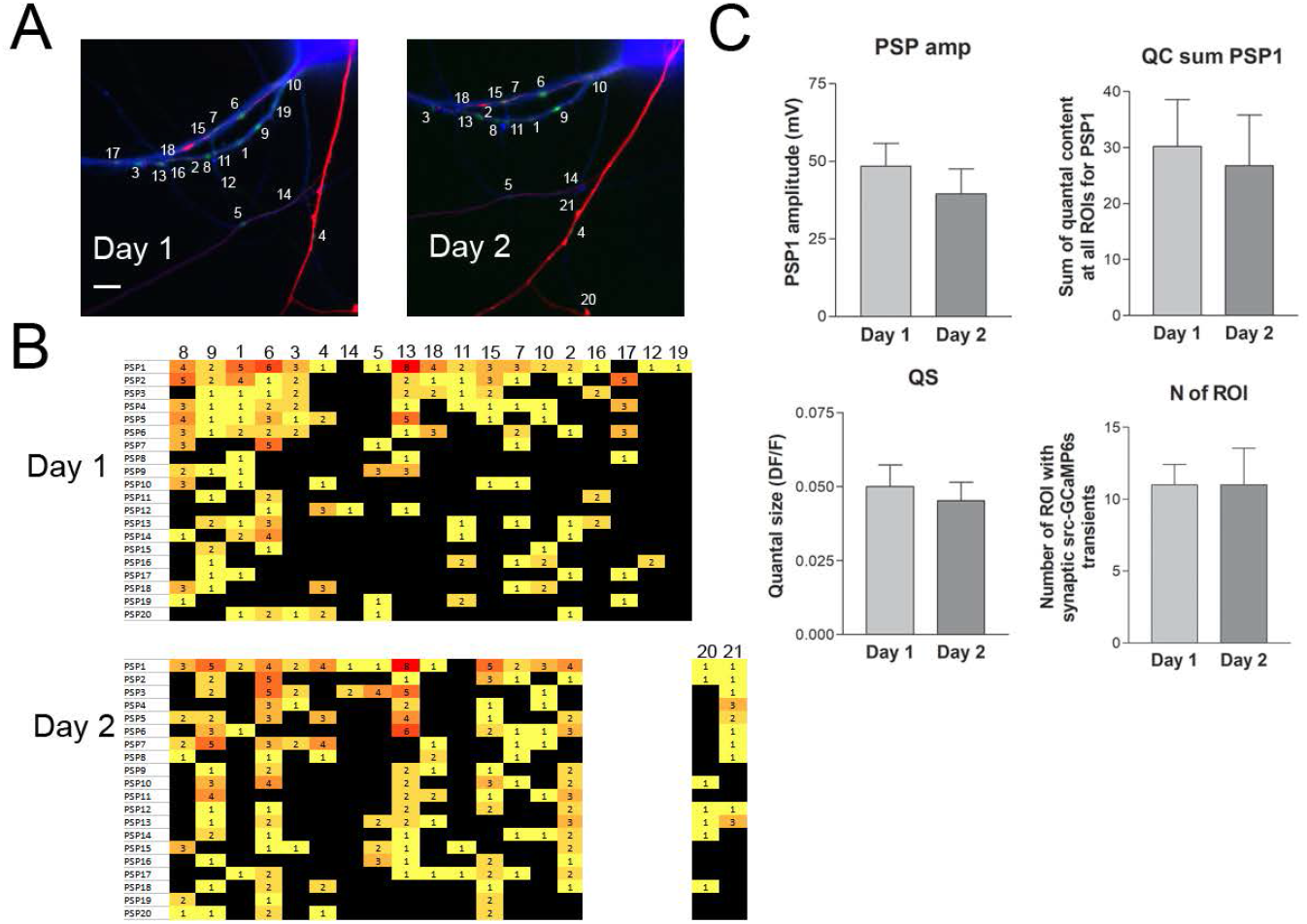
Tracking synapses over multiple days. A) Presynaptic RFP (red), postsynaptic src-GCaMP6s (blue), and the sum of all the single stimulus images (each generated with frame 11-frame 9 as in Fig.1D) compressed into a single image marking the location of the synapses (green). ROIs highlighted are selected as described above, and a blind judge categorizes each ROI on subsequent days as either an ROI identity from a preceding day, a new ROI, an ROI that has disappeared, or an ROI that may have appeared or disappeared from the field of view. This judges table is then used to track the ROIs over multiple days. Some ROIs such as 17 and 20 may have left or entered the field of view respectively, while ROIs like 12, 16 and 19 were not observed on the second day and others like 21 appear on the second day. B) The quantal content heat maps generated as described in figure 5, reorganized with the blind judges table to place ROIs judged to be the same on multiple days in the same column. Data from the same synaptic connection imaged in A. C) Summary data from five synaptic connections followed over 2 days with at least 5 ROI. Mean data for the amplitude of PSP1 (in mV), the average sum of quantal content for PSP1 and the average quantal size both measured as described in figure 4, and the number of ROIs with synaptic transients. Paired t-tests find no significant difference between any of the measures on day 1 and day 2, P>0.05, n=5.

## Discussion

### Src-GCamp6S can be used to monitor synaptic locations in sensory-motor neuron cultures

A functional synapse requires presynaptic release and postsynaptic receptors. Unlike measurements such as synaptophluorins or phluorin-tagged glutamate receptors, EPSCaTs actually require a functional synapse to be measured. We are able to largely eliminate issues with other calcium transients by locking our measurements to presynaptic action potentials and strict requirements for the timing of the rise of the EPSCaT. Moreover, since we summate over 20 action potentials, we should measure synapses with a probability of release over 0.05. The similar variance of uniquantal EPSCaTs and miniature PSPs suggests that the large EPSCaT fluctuations at some ROIs are multiquantal fluctuations, and that individual sites with expected single quantal fluctuations are measurable, an important benchmark for the technique as otherwise measurement would bias to multiquantal sites if uniquantal sites slip below detection.

### Multi-quantal ROIs

Our results show that even with the improved resolution of the membrane delimited Src-GCAMP6s, many EPSCaTs still appear to be multiquantal. While we cannot rule out the potential contribution of intrasite multivesicular release (Kusick et al. 2020), it is likely this largely results from multiple independent synapses existing in close proximity and such arrangements would not be resolvable with the methods used here. The propensity for sites to occur in clusters that are resolvable as in figure 4 further supports the idea that the ROIs with multiquantal EPSCaTs are a tight cluster of individual synapses.

It is interesting that a about half of the quantal content measured by the EPSCaTs comes from these apparent multiquantal sites. If these represent clustered synapses, it will be interesting in the future to learn why synapses are clustered together in one small region of the possible contacts between the sensory and motor neuron. We did not observe any obvious way of predetermining where this was in the synaptic arbor.

### Relationship between varicosities and synaptic sites

We find that only a minority of presynaptic varicosities have EPSCaTs apposed to them. These numbers are similar to those found with EPSCaTs measured with a calcium dye (Malkinson and Spira 2010), the number of presynaptic varicosities that have active release sites as measured with synaptophluorin (Kim et al. 2003), and the number of presynaptic varicosities in the animal that have active zones (Bailey and Chen 1983). For synaptophluorin and presynaptic varicosities in the animal, stimulation that leads to facilitation/sensitization increases the percentage of presynaptic varicosities with release sites and with active zones (Bailey and Chen 1983; Kim et al. 2003). We have not yet examined our results after inducing plasticity, and thus it is likely that the percentage of varicosities that have synapses may increase under these conditions. The more striking finding is the large number of EPSCaTs that occur at sites not adjoining presynaptic varicosities. This is approximately double the number previously measured with EPSCaTs in L7 neurons (Malkinson and Spira 2010). This could be due to differences between L7 and LFS neurons, differences in the details of the culture conditions, the focus on varicosity rich regions in the previous study, or increased sensitivity due to the use of src-GCAMP6s as opposed to a calcium dye. Earlier studies in L7 with EM to examine where active zones were located in cultures only focused on synaptic varicosities (Glanzman et al. 1989). Experiments in the animal do not explicitly discuss whether or not active zones are found outside varicosities (Bailey and Chen 1983). Another quality found with this resolution of the synapse was the propensity for the underlying elements to cluster together, further reducing the spatial location of synaptic activity to very small areas within the pre and postsynaptic contact. Our results strongly demonstrate, that is some circumstances, synapses in Aplysia sensory-motor neuron cultures, are not always primarily located at presynaptic varicosities and thus, the use of varicosities as a surrogate for synapses has caveats.

### The use of EPSCaTs to follow synapses over time

The stability of synapses is an important challenge for models where changes in synaptic strength underlie long-lasting memory. The ability to follow individual synaptic elements over time will allow us to determine what molecular elements may determine the stability of individual synapses and if certain synapses are particularly sensitive to memory disrupting agents such as persistent kinase inhibitors. The stability of synaptic efficacy over two days indicates that the sensor and the measurements do not affect the synapse, whereas the stability of the quantal size estimate demonstrates that the src-GCaMP6s synaptic subpopulation is consistent over many days allowing for the faithful detection of synaptic transmission.

## Methods

### Src-GCaMP6s generation

The generation of pNEXp-APsrc-gCAMP6s (Src-GCaMP6s) with the firsts 34 amino acids of Aplysia Src (NP_001191639.1) inserted in front of GCaMP6s was previously described (Farah et al. 2019). The first 34 amino acids of *Aplysia* Src contains conserved myristoylation and palmitylation sites that lead to membrane insertion.

### Cell culture

Aplysia were obtained from the National Resource for Aplysia at the University of Miami, and maintained in holding tanks until use. Aplysia ganglia were digested in dispase II and pleural sensory neurons were paired in isolation with LFS motor neurons in an isotonic L-15 based culture media containing 30% hemolymph and supplemented with L-glutamine in glass bottomed culture dishes and left for two days. On day three in culture, the nuclei of the neurons were injected with pNEX3 expression plasmids (Kaang et al. 1992) containing RFP (presynaptic neuron) and src-GCaMP6s (postsynaptic neuron). An additional 24h was allowed for plasmid expression before measuring synaptic transmission. For multi-day recordings, a portion of the culture media is retained so that when day 1 recording are finished the recording saline can be immediately replaced with fresh culture media (freshly supplemented with L-glutamine).

#### Electrophyisology

On day four, cell pairs expressing RFP presynaptically and src-GCaMP6s postsynaptically are examined in a high calcium recording saline [in mM NaCl 460, CaCl2 55, MgCl2 10, KCl 10, HEPES 10, D-glucose 10, pH 7.6]. Sharp electrodes (~15MO, back-filled with saturated potassium acetate) were used to control and hold membrane potential at −80mV and generate action potentials with an Axoclamp 900 and Digidata 1440 (Molecular Devices). Spontaneous miniature ‘mini’ PSPs were measured for a couple minutes prior to evoked presynaptic stimuli. Minis were identified and were measured with Clampfit (Molecular Devices) template event detection, using a template prepared previously from over one hundred minis at Aplysia pleural sensory neuron to LFS motor neuron synapses. Presynaptic action potentials are generated with a superthreshold depolarizing pulse 10ms in duration 0.56s into to the acquisition period initiated with activation of the imaging software. Postsynaptic potential (PSP) amplitude was measured with Clampfit.

### Fluorescence imaging

Cell pairs were imaged with a Zeiss Axio Observer D1, with a EC Plan Neofluar 1.3NA 40x oil coupled lens and a QuantEM: 512SC EM-CCD camera (Photometrics). Fluorescence excitation and emission achieved with an X-cite Expo arc, through Ziess RFP 20 and Ziess GFP 38 filter cubes. For each stimulus, 45 frames are acquired at 16.7 frames/second, with a constant 25ms exposure time at all trials, adjusting for baseline intensity differences between trials with camera gain. Rapid image acquisition was achieved with a Zeiss SVB-1 microscope signal distribution box and Axiovision 4.8 software using the fast acquisition application (Zeiss). Initiation of the Axiovision image acquisition electronically triggers the electrophysiological recording and therefore the stimulus to ensure that the presynaptic action potential is evoked at the same time in each 45 frame acquisition period, usually during frame 10 (slight differences in the membrane properties between the different sensory neurons from trial to trial vary the precise timing of action potential generation between cell pairs).

### ROI selection

All image analysis was done with Image J (Schneider et al. 2012) and Excel or Calc spreadsheets (Microsoft, LibreOffice). Subtraction of frame 9 from frame 11 generates an image of the src-GCaMP6s transients occurring during the stimulus. This image is then smoothed (each pixel converted to average of its 3×3 neighborhood) to aid in generation of clean ROI borders, and ROIs selected with the wand tool to draw an ROI boundary that best fits the shape of the transient. Once an ROI has been selected, the time-course of the fluorescence signal at the ROI is examined over the 45 frames to ensure that it did not initiate before frame 10 and that the kinetics of the transient are consistent with measurement of a synaptic current (i.e. rise to a clear peak and slower decay than rise). Once an ROI is selected at a stimulus it cannot be removed with selection of further ROIs for subsequent stimuli within a trial. ROIs chosen in a previously analyzed stimulus within a trial can only be further divided into a cluster of smaller ROIs with analysis of subsequent stimuli. To aid in separation of closely spaced ROIs, ROI selection is done from PSP20 (stimulus 20) backwards to PSP1, as the first stimulus has the most amount of synaptic activity, blurring closely spaced ROI boundaries. Following creation of the full ROI set for a particular synaptic connection, fluorescence intensity is measured for all stimuli using this full ROI set.

### Fluorescence intensity measurements

Fluorescence intensity measurements for all twenty stimuli at the full ROI set for a particular synaptic connection were analyzed in excel, where data were first normalized using frames 1-9 at each ROI and for each stimulus averaged to give Fo. All fluorescence intensity measurements were normalized as (Frame X-Fo) / Fo or DF/Fo. The standard deviation of frames 1-9 is used to measure the noise band before the stimulus for each ROI and every stimulus to define a signal detection limit. For a src-GCaMP6s transient to be considered a synaptic signal and further examined with quantal analysis, the DF/F at frame 11 needs to be greater than 3 times the standard deviation of frames 1-9. Visualization of this transient is achieved by subtracting the intensity values for each pixel in the image of frame 9 from the image of frame 11, creating an image of the src-GCaMP6s transient during the stimulus (see Fig. 1B). To visualize the transients at all the synapses participating over the twenty stimuli, the pixel intensity values of each of the above generated single stimulus transients are summed over all 20 stimuli to produce a single summated image of all the synaptic transients in a trial (see Fig. 1C). Note that in this image the intensity of an ROI is proportional to the quantal content at the site and non-synaptic calcium events that occur during any of the stimuli can enter this image, see below. Not all ROI chosen have a synaptic transient either because no transient is 3xSD 1-9, or all transients are eliminated due to non-synaptic calcium events. Thus, in the manuscript we distinguish between the ‘ROI set’ and the ‘ROI set of synaptic transients’.

If the frame preceding the stimulus has a fluorescent intensity change greater than the noise band for a stimulus at an ROI (DF/F frame9 > 3xSD frame1-8), the transient is considered not from calcium entering synaptic glutamate receptors (non-synaptic) as it initiated prior to the stimulus and is thus removed from subsequent analysis. So that the excitatory postsynaptic calcium transient (EPSCaT) for a specific stimulus is measured from the sum of DF/F changes at all ROIs except those with a non-synaptic transient preceding the stimulus as above. Examination of eighteen synaptic connections with at least five ROIs finds the rate of these non-synaptic calcium events can vary from trial to trial and from recording period to recording period ranging from none observed, up to ~4%, averaging 0.9±0.2%. Similarly, it can be estimated that a similar percentage of non-synaptic events initiate at frame 10 or 11 and thus coincidentally with the synaptic transients. Thus, about ~1% of the transients at ROIs judged as synaptic events are likely false positive non-synaptic transients. Such events can enter the images generated for the src-GCaMP6s transients as non-synaptic events, so not all fluorescence observed in these calculated images represents synaptic transients.

### Quantal analysis

Quantal size estimates were made at each ROI using the variance in the successful synaptic transients at each specific ROI. The estimates were made automatically with excel spreadsheets using a cluster analysis. At each ROI, the minimum, median, and maximum src-GCaMP6s synaptic transient is found. All events are then put in a cluster according to proximity to either the minimum, median, or maximum. The similarity of the cluster means is used to calculate quantal size, where if the mean of the minimum cluster is greater than 50% of the mean of the maximum, all events are considered uniquantal and averaged. If not, but the mean of the minimum is greater than 50% of the mean of the median, then all events in the minimum and median clusters are averaged for quantal size. Otherwise, only the mean of the minimum events is used to estimate quantal size. The 50% threshold used to find similarity between clusters was chosen as quantal increments would lead to transients twice the amplitude of the minimum with two quanta released, thus the minimum would be half or less with multiquantal synaptic transmission at an ROI.

### Statistical analysis

Statistical analyses were performed in Graph Pad Prism and Microsoft Excel. The larger data set for these experiments is from 13 synaptic pairs with at least 5 ROI with at least one transient judged to be synaptic, 1 trial was excluded as an ROI did not have a failure (a feature excluding quantal analysis) and 2 trials were excluded for a lack of correlation between PSP amplitude and the EPSCaT amplitude (as presented in Fig. 1, both of which appear to lose correlation due to large amounts of non-synaptic calcium transients). Five trials were examined on two consecutive days. In some cases (for example, data on non-synaptic transients) we use data from both days resulting in 18 synaptic pairs examined). Correlation of PSP amplitude to EPSCaT transient intensity was measured with a Pearson r calculation in Excel and a test statistic generated with the data set compared to a Students T distribution to generate a P value. All values are means ± standard error of the mean unless specifically stated otherwise.

## Acknowledgements

This work was supported by Canadian Institute of Health Research (CIHR) grant 340328 and an NSERC Discovery grant to WSS. WSS is a James McGill Professor.

